## Supplementary material for "Multiple Notch ligands in the synchronization of the segmentation clock": Suplemental Material

### I. REACTION KINETICS AND MODEL FORMULATION

**Regulatory function.** This work focuses on the roles of different components of the Notch signaling network, so we take a parsimonious approach to describe the core oscillator. We consider a single Her protein that inhibits its own production with a time delay accounting for synthesis time. The single variable $H(t)$ represents the concentration of this protein at time $t$. Besides self inhibition, the synthesis rate of Her protein is positively regulated by Notch signaling components. The Notch intracellular domain (NICD) is released inside the cell upon binding of ligands from neighboring cells to the Notch receptor. We denote the concentration of this signaling component by $S(t)$.

Here, we derive a function to describe the regulatory effects of $H$ and $S$ on the synthesis rate of $H$. We will first consider the dynamics and steady state of the promoter and the binding factors. The promoter architecture of *her* genes may be complex, including about 12 binding sites for the Her proteins [1]. Besides, it is thought that Her proteins bind as dimers to their own promoters [1]. It has been shown that multiple binding sites effectively increase the nonlinearity of regulatory functions [2–4]. Similarly, dimerization can be effectively described in an adiabatic approximation by a Hill exponent of 2 [1]. Thus, here we assume that there is only one binding site for Her proteins for simplicity, and then include the effects of multiplicity in an effective Hill exponent [2]. We further assume that NICD binds as a single molecule to the activating complex at the Her promoter.

With these assumptions, the promoter can be in any of four states, depending on whether none, one, or both of the binding sites are occupied. We represent the promoter state as $P_{\mu\nu}$, where $\mu$ is the state of the Her binding site and $\nu$ is the state of the NICD binding site, with $\mu, \nu = 0$ indicating an empty site and $\mu, \nu = 1$ an occupied site. The different promoter states are connected in the scheme of binding reactions

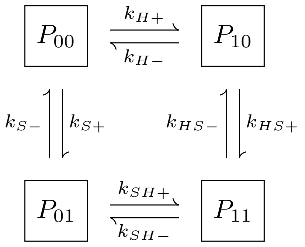


where the + and − signs label binding and unbinding rates respectively. The values of $k_{HS\pm}$ and $k_{SH\pm}$ depend on how the two regulatory components interact. For instance, setting them both to zero, $k_{HS+} = k_{SH+} = 0$ describes exclusive competitive binding, where the doubly bound promoter state is not accessible [5]. Alternatively, assuming the binding rates to be independent of the promoter state for both components, $k_{SH\pm} = k_{H\pm}$ and $k_{HS\pm} = k_{S\pm}$, describes a situation with no interaction at the binding sites of regulatory components, which we term dual binding model. Further choices describe models for different ligand interactions. Motivated by the multiple binding sites accessible to Her transcription factors, here we focus on the dual binding model. The reactions involved in this dual binding scenario are

$$\mathrm{P}_{00} + \mathrm{H} \underset{\mathrm{k_{H-}}}{\overset{\mathrm{k_{H+}}}{\rightleftharpoons}} \mathrm{P}_{10}\,, \qquad \mathrm{P}_{10} + \mathrm{S} \underset{\mathrm{k_{S-}}}{\overset{\mathrm{k_{S+}}}{\rightleftharpoons}} \mathrm{P}_{11}\,, \tag{1}$$

$$\mathrm{P}_{00} + \mathrm{S} \underset{\mathrm{k_{S-}}}{\overset{\mathrm{k_{S+}}}{\rightleftharpoons}} \mathrm{P}_{01}\,, \qquad \mathrm{P}_{01} + \mathrm{H} \underset{\mathrm{k_{H-}}}{\overset{\mathrm{k_{H+}}}{\rightleftharpoons}} \mathrm{P}_{11}\,. \tag{2}$$

From these reactions we can write the dynamics for the four states of the promoter

$$\dot{P}_{00} = k_{H-}P_{10} + k_{S-}P_{01} - k_{H+}HP_{00} - k_{S+}SP_{00}\,, \tag{3}$$

$$\dot{P}_{10} = k_{H+}HP_{00} + k_{S-}P_{11} - k_{H-}P_{10} - k_{S+}SP_{10}\,, \tag{4}$$

$$\dot{P}_{01} = k_{S+}SP_{00} + k_{H-}P_{11} - k_{S-}P_{01} - k_{H+}HP_{01}\,, \tag{5}$$

$$\dot{P}_{11} = k_{H+}HP_{01} + k_{S+}SP_{10} - k_{H-}P_{11} - k_{S-}P_{11}\,, \tag{6}$$

together with the conservation law

$$P_{00} + P_{10} + P_{01} + P_{11} = P_T\,. \tag{7}$$

We assume that promoter binding and unbinding reactions are much faster than the dynamics of $H$ and $S$. With this assumption, the promoter state is in equilibrium for given concentrations of $H$ and $S$ and we set $\dot{P}_{\mu\nu} = 0\ \forall \mu, \nu$. We can solve the resulting algebraic equations to obtain the quasi-steady state promoter occupancy $P_{\mu\nu}(H, S)$,

$$P_{00}(H,S) = \frac{k_{H-}k_{S-}P_T}{(k_{H-} + k_{H+}H)\,(k_{S-} + k_{S+}S)}\,, \tag{8}$$

$$P_{10}(H,S) = \frac{k_{S-}k_{H+}P_T H}{(k_{H-} + k_{H+}H)\,(k_{S-} + k_{S+}S)}\,, \tag{9}$$

$$P_{01}(H,S) = \frac{k_{H-}k_{S+}P_T S}{(k_{H-} + k_{H+}H)\,(k_{S-} + k_{S+}S)}\,, \tag{10}$$

$$P_{11}(H,S) = \frac{k_{H+}k_{S+}P_T HS}{(k_{H-} + k_{H+}H)\,(k_{S-} + k_{S+}S)}\,. \tag{11}$$

Next, we consider how promoter occupancy determines transcription rates. We introduce the regulatory function $f(H, S)$ that modulates the synthesis rate of $H$

$$f(H,S) = \sum_{\mu,\nu=0,1} a_{\mu\nu}P_{\mu\nu}(H,S)\,, \tag{12}$$

where each promoter state $P_{\mu\nu}$ has an associated transcription rate $a_{\mu\nu}$. Here assume that binding of Her to the promoter fully represses synthesis independently of the binding of NICD, $a_{10} = a_{11} = 0$. We set a basal transcription rate $a_{00} = b$ and an activated state to a higher rate $a_{01} = a > b$, resulting in

$$f(H,S) = P_T \frac{k_{H-}(bk_{S-} + ak_{S+}S)}{(k_{H-} + k_{H+}H)(k_{S-} + k_{S+}S)} \,. \qquad (13)$$

Introducing concentration scales

$$H_0 \equiv \frac{k_{H-}}{k_{H+}} \quad \text{and} \quad S_0 \equiv \frac{k_{S-}}{k_{S+}} \,, \qquad (14)$$

we arrive at

$$f(H,S) = \frac{b_H}{1+\frac{H}{H_0}} \frac{1+\frac{a_H}{b_H}\frac{S}{S_0}}{1+\frac{S}{S_0}} \,, \qquad (15)$$

where we defined $a_H \equiv P_T a$ and $b_H \equiv P_T b$.

This expression provides the structure of the regulatory function. As we described above, Her proteins form dimers and these dimers bind to multiple binding sites on the promoter. Similarly, the NICD binds DNA through a complex that may also introduce additional nonlinearities. These effects can be encompassed in effective Hill exponents $h_H$ and $h_S$,

$$f(H,S) = b_H \frac{1}{1+\left(\frac{H}{H_0}\right)^{h_H}} \frac{1+\frac{a_H}{b_H}\left(\frac{S}{S_0}\right)^{h_S}}{1+\left(\frac{S}{S_0}\right)^{h_S}} \,. \qquad (16)$$

The regulatory function separates into factors,

$$f_-(H) = \frac{1}{1+\left(\frac{H}{H_0}\right)^{h_H}} \quad \text{and} \quad f_+(S) = \frac{1+\frac{a_H}{b_H}\left(\frac{S}{S_0}\right)^{h_S}}{1+\left(\frac{S}{S_0}\right)^{h_S}} \,, \qquad (17)$$

such that

$$f(H,S) = b_H f_-(H) f_+(S) \,, \qquad (18)$$

reflecting the independent binding assumption, where binding to one site is independent of the state of the other.

Next, to complete the formulation of the model, we describe the different reactions involved, from synthesis to degradation, ligand dimerization, and Notch binding by different components.

**Synthesis.** Her synthesis proceeds at a basal rate $b_H$, modulated by the regulatory function Eq. (16),

$$\dot{H}_i(t) = b_H \frac{1}{1+\left(\frac{H_i(t-\tau_i)}{H_0}\right)^{h_H}} \frac{1+\frac{a_H}{b_H}\left(\frac{S_i(t-\tau_i)}{S_0}\right)^{h_S}}{1+\left(\frac{S_i(t-\tau_i)}{S_0}\right)^{h_S}} + \cdots \,, \qquad (19)$$

where $i = 1, \ldots, N_c$ is the cell label and $\tau_i$ is an explicit synthesis delay accounting for the multiple steps in the negative feedback of $H$ [6, 7]. Since a Her molecule takes a time $\tau_i$ to be synthesized in cell $i$, the synthesis rate at time $t$ depends on Her and NICD concentrations at a previous time $t - \tau_i$, when synthesis was starting. This delay is different for each cell, introducing a variability in the period of the cell population as described below.

To describe DeltaC synthesis inhibition by Her, we follow a similar strategy as above. Considering the reaction kinetics for the binding of Her to the DeltaC promoter,

$$\dot{C}_i(t) = b_C \frac{1}{1+\left(\frac{H_i(t-\tau_C)}{H_{0C}}\right)^{h_C}} + \cdots \,, \qquad (20)$$

where $b_C$ is the basal rate, $\tau_C$ is the DeltaC synthesis delay, $H_{0C}$ is the threshold for Her mediated inhibition of DeltaC synthesis and $h_C$ is an effective Hill exponent describing nonlinear effects in the inhibition. Finally, we set constant synthesis rates for DeltaD and Notch, since they are not regulated by other components from the oscillator or the coupling network, resulting in constant terms

$$\dot{D}_i(t) = b_D + \cdots \qquad \text{and} \qquad \dot{N}_i(t) = b_N + \cdots \,. \qquad (21)$$

**Degradation.** We assume decay of all components according to the reactions

$$\mathrm{X} \xrightarrow{d_X} \varnothing \,, \qquad (22)$$

resulting in a linear decay term

$$\dot{X}_i(t) = -d_X X_i(t) + \cdots \qquad (23)$$

for all variables $X = H, C, D, E, F, G, N, S$, where $d_X$ is the decay rate for the variable $X$.

**Ligands dimerization.** Notch ligands DeltaC and DeltaD can form an heterodimer and both homodimers, described in the reactions

$$\mathrm{C}_i + \mathrm{D}_i \underset{\mathrm{k_{CD}}^-}{\overset{\mathrm{k_{CD}}^+}{\rightleftharpoons}} \mathrm{C{:}D}_i \,, \qquad (24)$$

$$2\,\mathrm{C}_i \underset{\mathrm{k_{CC}}^-}{\overset{\mathrm{k_{CC}}^+}{\rightleftharpoons}} \mathrm{C{:}C}_i \,, \qquad (25)$$

$$2\,\mathrm{D}_i \underset{\mathrm{k_{DD}}^-}{\overset{\mathrm{k_{DD}}^+}{\rightleftharpoons}} \mathrm{D{:}D}_i \,. \qquad (26)$$

We introduce the notation for dimers concentrations, [DeltaC:DeltaD] $\equiv E$, [DeltaC:DeltaC] $\equiv F$, [DeltaD:DeltaD] $\equiv G$, and relabel the corresponding rates as $k_E^{\pm} \equiv k_{CD}^{\pm}$, $k_F^{\pm} \equiv k_{CC}^{\pm}$, and $k_G^{\pm} \equiv k_{DD}^{\pm}$. Each reaction contributes terms to the components involved. For example, Eq. (25) will contribute terms to the equations for the DeltaC monomer and the DeltaC:DeltaC homodimer,

$$\dot{C}_i(t) = -2k_F^+ C_i^2 + 2k_F^- F_i + \cdots \,, \qquad (27)$$

$$\dot{F}_i(t) = +k_F^+ C_i^2 - k_F^- F_i + \cdots \,. \qquad (28)$$

**Notch binding.** Ligands from a neighboring cell, in the form of monomers or dimers, may bind to a Notch receptor. Once Notch is bound by a ligand, its intracellular domain is released into the cell and the complex formed by the remaining extracellular domain and the bound ligand is internalized into the other cell. Here we assume that the ligand is degraded after the interaction with a receptor. Since the receptor is cleaved in the process, we describe this as an irreversible reaction,

$$\mathrm{X}_i + \mathrm{N}_j \xrightarrow{\mathrm{k_{NX}}} \mathrm{S}_j \qquad , \qquad \mathrm{X} = \mathrm{C,\ D,\ E,\ F,\ G}\,, \tag{29}$$

where $k_{NX}$ is the binding rate between Notch and a ligand $X$, and $i, j$ label the interacting cells.

**Average neighbor coupling.** These Notch binding reactions give rise to the coupling terms in the dynamics of $X$

$$\dot{X}_i(t) = -\sum_{j\in\mathcal{V}_i} k_{NX} X_i N_j r_{ij} + \cdots\,, \tag{30}$$

where the sum is over neighbors $\mathcal{V}_i$ of cell $i$. The contribution from each neighbor $j$ is weighted by the fraction $r_{ij}$ of the total surface of cell $i$ in contact with cell $j$,

$$r_{ij} = \frac{A_{ij}}{A_T}\,, \tag{31}$$

where $A_{ij}$ is the contact area between the two neighbors and $A_T$ is the total area of a cell, assumed to be the same for all cells. Assuming that the total cell $i$ area is shared equally with all $|\mathcal{V}_i|$ neighbors, $A_{ij} = A_T/|\mathcal{V}_i|$, and

$$r_{ij} = \frac{1}{|\mathcal{V}_i|}. \tag{32}$$

With this assumption, cell $i$ is coupled to the average of Notch concentrations in neighboring cells,

$$\dot{X}_i(t) = -k_{NX} X_i \frac{1}{|\mathcal{V}_i|}\sum_{j\in\mathcal{V}_i} N_j + \cdots\,. \tag{33}$$

Similar average couplings arise in the equations for $N$ and $S$,

$$\dot{N}_i(t) = -k_{NX} N_i \frac{1}{|\mathcal{V}_i|}\sum_{j\in\mathcal{V}_i} X_j + \cdots\,, \tag{34}$$

$$\dot{S}_i(t) = +k_{NX} N_i \frac{1}{|\mathcal{V}_i|}\sum_{j\in\mathcal{V}_i} X_j + \cdots\,. \tag{35}$$

**Mean field.** In this work we focus on the tailbud, a posterior region of the segmentation clock characterized by synchronized oscillations in a population of very mobile cells [8]. This cell mobility causes a continuous neighbor exchange, and it has been shown that the resulting dynamics can be effectively described as an all-to-all coupling, or mean field [9]. Thus, we write the average terms in Eqs. (33)-(35) for the ligands and the Notch receptor as mean field variables

$$\bar{X} = \frac{1}{N_c}\sum_{i=1}^{N_c} X_i \qquad , \qquad X = C, D, E, F, G, N. \tag{36}$$

With these definitions, we can write the gain and loss terms that reactions in Eq. (29) contribute to different components. For example, the heterodimer E participates in the contributions

$$\dot{S}_i(t) = +k_{NE}\bar{E}N_i + \cdots\,, \tag{37}$$

$$\dot{E}_i(t) = -k_{NE}E_i\bar{N} + \cdots\,, \tag{38}$$

$$\dot{N}_i(t) = -k_{NE}\bar{E}N_i + \cdots\,. \tag{39}$$

**Full model.** Putting all contributions together, we have eight coupled delayed differential equations for each cell $i$,

$$\dot{H}_i = -d_H H_i + b_H \frac{1}{1+\left(\frac{H_i(t-\tau_i)}{H_0}\right)^{h_H}} \frac{1+\frac{a_H}{b_H}\left(\frac{S_i(t-\tau_i)}{S_0}\right)^{h_S}}{1+\left(\frac{S_i(t-\tau_i)}{S_0}\right)^{h_S}} \tag{40}$$

$$\dot{C}_i = -d_C C_i + b_C \frac{1}{1+\left(\frac{H_i(t-\tau_C)}{H_{0C}}\right)^{h_C}} - k_E^+ C_i D_i + k_E^- E_i - 2k_F^+ C_i^2 + 2k_F^- F_i - k_{CN} C_i \bar{N} \tag{41}$$

$$\dot{D}_i = -d_D D_i + b_D + k_E^- E_i - k_E^+ C_i D_i - 2k_G^+ D_i^2 + 2k_G^- G_i - k_{DN} D_i \bar{N} \tag{42}$$

$$\dot{E}_i = -d_E E_i - k_E^- E_i + k_E^+ C_i D_i - k_{EN} E_i \bar{N} \tag{43}$$

$$\dot{F}_i = -d_F F_i - k_F^- F_i + k_F^+ C_i^2 - k_{FN} F_i \bar{N} \tag{44}$$

$$\dot{G}_i = -d_G G_i - k_G^- G_i + k_G^+ D_i^2 - k_{GN} G_i \bar{N} \tag{45}$$

$$\dot{N}_i = -d_N N_i + b_N - k_{CN}\bar{C}N_i - k_{DN}\bar{D}N_i - k_{EN}\bar{E}N_i - k_{FN}\bar{F}N_i - k_{GN}\bar{G}N_i \tag{46}$$

$$\dot{S}_i = -d_S S_i + k_{CN}\bar{C}N_i + k_{DN}\bar{D}N_i + k_{EN}\bar{E}N_i + k_{FN}\bar{F}N_i + k_{GN}\bar{G}N_i \tag{47}$$

where we omit the time dependence for notational simplicity except in delayed contributions. This full model is a superset of all the special cases considered in the main text, including all possible dimers and Notch binding species. The scenarios considered in the main text can be obtained by an adequate choice of parameters. For instance, choosing the binding rates $k_{CN} = k_{DN} = 0$ gives us the dimer binding scenario of the main text, while choosing all dimerization constants $k_X^{\pm} = 0$ and the corresponding Notch coupling constants $k_{NX} = 0$ for $X = E, F, G$ results in the monomer binding scenario.

**Period variability.** The autonomous period of the core oscillator $i$ depends primarily on the value of the synthesis delay $\tau_i$ in the negative feedback, and the half life of the protein $d_H^{-1}$, with second order corrections depending mainly on the synthesis rate $b_H$ [10]. To introduce variability in the autonomous periods, we sample the delays $\tau_i$ of individual oscillators from a normal distribution, with mean $\tau_H$ and variance $\sigma_\tau$.

### II. DIMENSIONLESS FORMULATION

Next, we introduce the dimensionless formulation of the model that we use in the main text, which provides a description that is independent of timescales and concentration scales. The equations (40)-(47) have units of concentration over time. We choose the concentration scale $H_0$ and the timescale $d_H^{-1}$, and render the equations dimensionless multiplying all by the same factor $(d_H H_0)^{-1}$. Introducing dimensionless variables

$$x_i \equiv \frac{X_i}{H_0} \quad , \quad X = H, C, D, E, F, G, N, S \,, \tag{48}$$

and dimensionless time

$$\tilde{t} \equiv t d_H \,, \tag{49}$$

the time derivatives transform as

$$\frac{\dot{X}}{d_H H_0} = \frac{dx}{d\tilde{t}} \equiv x' \,, \tag{50}$$

where we defined the prime notation for the derivative with respect to $\tilde{t}$. The resulting dimensionless formulation is

$$h_i' = -h_i + \beta_h \frac{1}{\left[1 + h_i^{\eta_H}(\tilde{t} - \tilde{\tau}_i)\right]} \frac{1 + \alpha\left(\sigma s_i(\tilde{t} - \tilde{\tau}_i)\right)^{\eta_S}}{\left[1 + \left(\sigma s_i(\tilde{t} - \tilde{\tau}_i)\right)^{\eta_S}\right]} \tag{51}$$

$$c_i' = -\delta_c c_i + \beta_c \frac{1}{1 + \left(\gamma h_i(\tilde{t} - \tilde{\tau}_C)\right)^{\eta_C}} + \lambda_E^- e_i - \lambda_E^+ c_i d_i + \lambda_F^- f_i - \lambda_F^+ c_i^2 - \kappa_C c_i \bar{n} \tag{52}$$

$$d_i' = -\delta_d d_i + \beta_d + \lambda_E^- e_i - \lambda_E^+ c_i d_i + \lambda_G^- g_i - \lambda_G^+ d_i^2 - \kappa_D d_i \bar{n} \tag{53}$$

$$e_i' = -\delta_e e_i - \lambda_E^- e_i + \lambda_E^+ c_i d_i - \kappa_E e_i \bar{n} \tag{54}$$

$$f_i' = -\delta_f f_i - \lambda_F^- f_i + \lambda_F^+ c_i^2 - \kappa_F f_i \bar{n} \tag{55}$$

$$g_i' = -\delta_G g_i - \lambda_G^- g_i + \lambda_G^+ d_i^2 - \kappa_G g_i \bar{n} \tag{56}$$

$$n_i' = -\delta_n n_i + \beta_n - n_i \sum_{c,d,e,f,g} \kappa_X \bar{x} \tag{57}$$

$$s_i' = -\delta_s s_i + n_i \sum_{x=c,d,e,f,g} \kappa_X \bar{x} \tag{58}$$

with dimensionless parameters

$$\delta_X \equiv \frac{d_X}{d_H} \quad , \quad \beta_X = \frac{b_X}{d_H H_0} \quad , \quad \kappa_X = \frac{k_{XN} H_0}{d_H} \,,$$

$$\sigma = \frac{H_0}{S_0} \quad , \quad \gamma \equiv \frac{H_0}{H_{0C}} \quad , \quad \alpha = \frac{a_H}{b_H} \,,$$

$$\lambda_E^+ \equiv \frac{H_0 k_E^+}{d_H} \quad , \quad \lambda_E^- \equiv \frac{k_E^-}{d_H} \,,$$

$$\lambda_F^+ \equiv \frac{2H_0 k_F^+}{d_H} \quad , \quad \lambda_F^- \equiv \frac{2k_F^-}{d_H} \,,$$

$$\lambda_G^+ \equiv \frac{2H_0 k_G^+}{d_H} \quad , \quad \lambda_G^- \equiv \frac{2k_G^-}{d_H} \,.$$

and renaming $h_X \to \eta_X$, $X = H, S, C$. For notational simplicity and readability, in the main text we drop the tildes from dimensionless time variables $\tilde{t} \to t$ and $\tilde{\tau}_X \to \tau_X$. We also drop the prime notation and use a dot to denote dimensionless time derivatives.

### III. PARAMETER VALUES

Next, we parametrize the model to describe both wild-type and mutant conditions. Experimental observations of the zebrafish segmentation clock, together with theoretical considerations, set constraints on parameter values. To parametrize the core oscillator we follow [10], comparing the mRNA and protein model

$$\dot{p}(t) = am(t-\tau_p) - bp(t)\,, \tag{59}$$

$$\dot{m}(t) = \frac{k}{1+\left(\frac{p(t-\tau_m)}{p_0}\right)^n} - cm(t)\,, \tag{60}$$

to the protein only oscillator from Eq. (51)

$$\dot{H}(t) = -d_H H(t) + \frac{b_H}{1+\left(\frac{H(t-\tau_H)}{H_0}\right)^{\eta_H}}\,. \tag{61}$$

Parameter values estimated for the mRNA and protein oscillator are [10]: $a = 4.5$ proteins per mRNA molecule, $c = 0.23$ molecules per minute, $b = 0.286$ molecules per minute estimated from experiments [11], $k = 33$ mRNA molecules per diploid cell per minute, and $p_0 = 40$ molecules. Synthesis delays in the model were later estimated from experiments [12], showing that her1 and her7 have very similar transcription times $\tau_{m1} \approx 10$ min and $\tau_{m7} \approx 9$ min, dominating over the protein translation times $\tau_{p1} \approx 2.8$ min and $\tau_{p7} \approx 1.7$ min.

Introducing a quasi-steady state approximation for the mRNA, $\dot{m} \approx 0$, we obtain

$$m(t) \approx \frac{k}{c}\frac{1}{1+\left(\frac{p(t-\tau_m)}{p_0}\right)^n}\,, \tag{62}$$

and replacing $m(t-\tau_p)$ into (59),

$$\dot{p}(t) = -bp(t) + \frac{ak}{c}\frac{1}{1+\left(\frac{p(t-\tau_m-\tau_p)}{p_0}\right)^n}\,. \tag{63}$$

Comparing Eqs. (61) and (63) we see that to obtain similar dynamics we should set $d_H \approx b$, $b_H \approx ak/c$, $\tau_H \approx \tau_m + \tau_p$, $\eta_H \approx n$ and $H_0 \approx p_0$. Since this comparison relies on the quasi-steady state approximation, a strict assignment of these values may produce differences in amplitude and period. In addition, the synthesis delay $\tau_H$ could be estimated differently from comparing the estimated period for both models,

$$T_{\text{Lewis}} \approx 2(\tau_p + \tau_m + 1/b + 1/c)\,, \tag{64}$$

$$T_H \approx 2(\tau_H + 1/d_H)\,. \tag{65}$$

For these periods to match, we would require $\tau_H \approx \tau_m + \tau_p + 1/c$.

For the protein only model, parameter values from [10] imply a Her protein degradation rate $d_H = 0.286$ min$^{-1}$, setting the timescale, and $H_0 = p_0 = 40$ molecules, corresponding to a concentration scale $[H_0] \approx 0.3$ nM for tailbud cells of 7.6 $\mu$m in diameter [8]. To obtain oscillations of similar amplitude to [10], we set $\beta_H = 28$ in the dimensionless model, corresponding to $b_H \approx 320$ proteins per minute. We set the dimensionless delay to $\tau_h = 4.2$, corresponding to a synthesis delay within a range of values obtained from averaging Her1 and Her7 data, $\tau_H = 14.7$. Hill exponents have an effective value that encompasses the effect of multiple macromolecular interactions in the regulation, such as dimerization [1], binding of complexes and multiple binding sites at the promoter [2, 3]. We expect exponent values to be larger than 2, so we set the conservative estimate $\eta = 2.5$. We will use these values for the core oscillator and determine the rest of parameters from further estimations and explorations.

The rest of the parameters concern signaling components, such as the ligands DeltaC and DeltaD, their dimerization, binding to Notch receptors and signal production and action on the oscillator. We set some of these parameters below and explore the effects of varying oth-

| parameter | dimers | monomers | description |
|---|---|---|---|
| $\delta_C$ | 1 | 1 | DC degradation rate |
| $\delta_D$ | 1 | 1 | DD degradation rate |
| $\delta_E$ | 1 | 1 | DC:DD degradation rate |
| $\delta_F$ | 1 | 1 | DC:DC degradation rate |
| $\delta_G$ | 1 | 1 | DD:DD degradation rate |
| $\delta_N$ | 1 | 1 | Notch degradation rate |
| $\delta_S$ | 1 | 1 | NICD degradation rate |
| $\beta_H$ | 28 | 28 | Her synthesis rate |
| $\beta_C$ | 26 | 39 | DC synthesis rate |
| $\beta_D$ | 17 | 20 | DD synthesis rate |
| $\beta_N$ | 24 | 30 | Notch synthesis rate |
| $\lambda_{E+}$ | 1 | – | DC:DD dimerization rate |
| $\lambda_{E-}$ | 0.1 | – | DC:DD dissociation rate |
| $\lambda_{F+}$ | 0.1 | – | DC:DC dimerization rate |
| $\lambda_{F-}$ | 0.1 | – | DC:DC dissociation rate |
| $\lambda_{G+}$ | 0.1 | – | DD:DD dimerization rate |
| $\lambda_{G-}$ | 0.1 | – | DD:DD dissociation rate |
| $\kappa_C$ | – | 0.02 | DC binding rate to Notch |
| $\kappa_D$ | – | 0.01 | DD binding rate to Notch |
| $\kappa_E$ | 0.2 | – | DC:DD binding rate to Notch |
| $\kappa_F$ | 0.02 | – | DC:DC binding rate to Notch |
| $\kappa_G$ | 0.009 | – | DD:DD binding rate to Notch |
| $\sigma$ | 0.1 | 0.1 | Signal activation threshold |
| $\gamma$ | 1 | 1 | DC inhibition threshold |
| $\eta_H$ | 2.5 | 7 | Her self-inhibition H.e. |
| $\eta_S$ | 2.5 | 7 | Her activation by signal H.e. |
| $\eta_C$ | 2.5 | 7 | DC inhibition by Her H.e. |
| $\alpha$ | 10 | 10 | Fold change in synthesis rate |
| $\tau_H$ | 4.2 | 4.2 | Mean Her synthesis delay |
| $\sigma_\tau/\tau_H$ | 0.03 | 0.03 | C.V. of Her synthesis delay |
| $\tau_C/\tau_H$ | 1.7 | 1.7 | Relative DC synthesis delay |

Table I. Parameter table used in synchronization maps in the main text. DC and DD stand for DeltaC and DeltaD. H.e. stands for Hill exponent. C.V. is coefficient of variation.

ers.

Next we consider signal reception parameters in Eq. (51). The dimensionless threshold for signal driven synthesis activation $\sigma$ controls how much signal is necessary to affect the core oscillator. We chose a value $\sigma = 0.1$, which corresponds to a weaker action than the inhibitor. For the signal to activate Her protein synthesis above basal levels we require a coupling strength $\alpha > 1$. Through exploration, we settled for a value $\alpha = 10$, which increases Her collective oscillation amplitude by a factor of $\sim 2$. The corresponding Hill exponent $\eta_S$ was set equal to $\eta_H$ for simplicity.

In the regulation of DeltaC synthesis by Her protein Eq. (52), we set the dimensionless threshold for synthesis repression $\gamma = 1$, assuming for simplicity that the same concentration of Her protein is required to inhibit both Her and DeltaC synthesis. Experimental observations in zebrafish showed that impairing Notch signaling results in longer collective period [13]. For reduced coupling to lengthen the period, effective coupling delays should be slightly below the period value [14, 15]. With this motivation we set the synthesis delay of DeltaC to $\tau_C = 1.7\tau_H$. The corresponding Hill exponent $\eta_C$ was set equal to $\eta_H$ for simplicity.

Given the lack of specific data for each component, we set all decay rates in the model to the same value $d_X = d_H$ with $X \in \{C, D, E, F, G, N, S\}$, for simplicity. So $\delta_X = 1$ for all $X$ in the dimensionless model. Motivated by experimental observations, we set the values of dimerization and binding rates to Notch of the two homodimers about an order or magnitude lower than those of the heterodimer [16]. Scans of these parameters revealed that dimer association rates $\lambda_X^+$ have maximal effective values: beyond some level, the size of the synchronization region in $\beta_C$ vs. $\beta_D$ space stops changing significantly. Dimer dissociation rates $\lambda_X^-$ were found to have small effects on the size and shape of the synchronization regions as long as they where within an order of magnitude of the corresponding $\lambda_X^+$, and were set to $\lambda_X^- = 0.1$.

The remaining parameters are the synthesis rates of coupling components and the binding rates of different components to Notch receptors. We vary synthesis rates $\beta_C$, $\beta_D$ and $\beta_N$ to construct synchronization maps in terms of these. Additionally, we vary the values of $\kappa_X$ to explore how these maps change.

### IV. SUPPLEMENTARY FIGURES

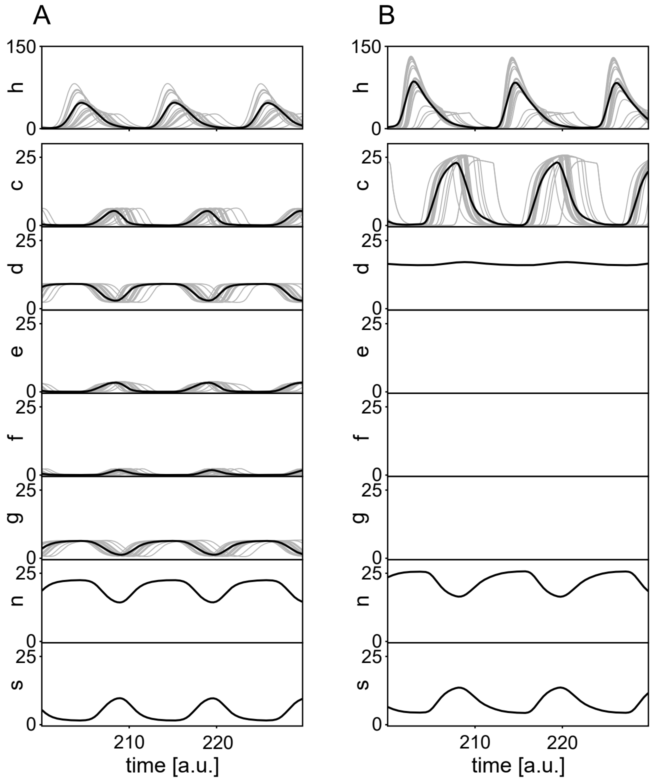


**Fig. S1**. Steady state individual oscillations $x_i(t)$ (grey lines, 25 out of 100 are displayed) and mean field $\bar{x}(t)$ (black line), for all variables $x = h, c, d, e, f, g, n, s$, in the (A) dimer scenario and (B) monomer scenario. $e(t)$, $f(t)$ and $g(t)$ are not defined in the monomer model, hence panels are empty. Vertical scales are the same except for $h(t)$. Parameters as in Table I.

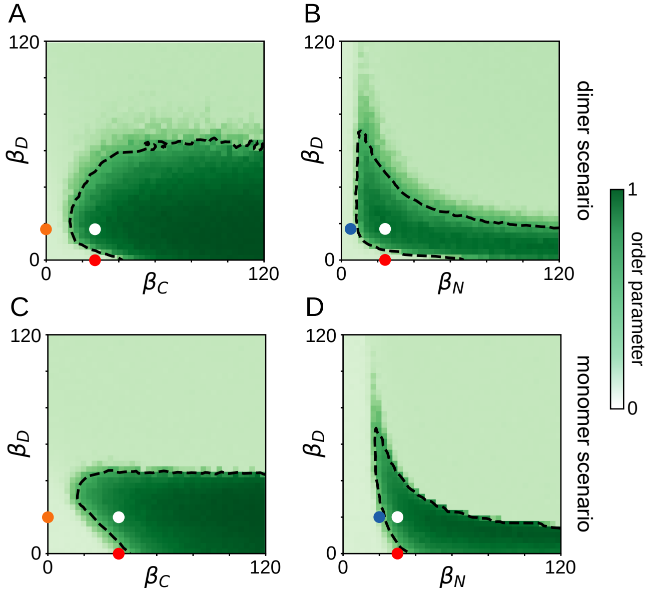


**Fig. S2**. Extended synchronization maps for wider ranges of the syntheses rates as indicated, for (A, B) the dimer scenario shown in Fig. 2 of the main text and (C, D) the monomer scenario shown in Fig. 5 of the main text. Colored dots indicate the parameter sets used in the desynchronization assays for each model. Dashed line and number of realizations as in Fig. 2C, D of the main text.

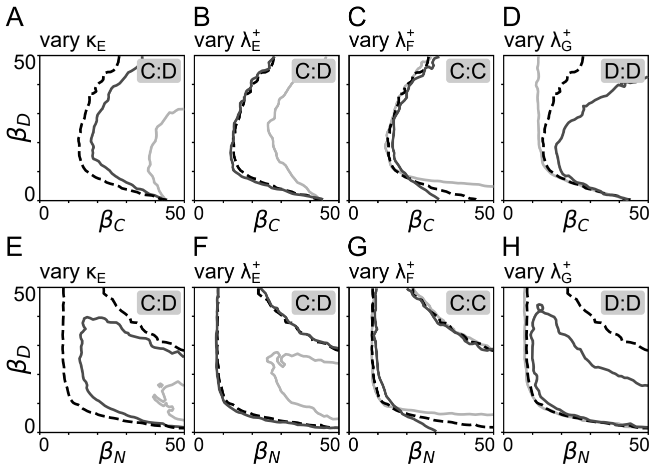


**Fig. S3**. Changes to the sync region boundary of the map (A-D) in Fig. 2C of the main text and (E-H) in Fig. 2D of the main text, as a function of different parameters of the model. (A, E) $\kappa_E = 0.02$ (light), 0.066 (dark), 0.2 (dashed black). (B, F) $\lambda_E^+ = 0.05$ (light), 1 (dashed black), 3.33 (dark). (C, G) $\lambda_F^+ = 0.006$ (light), 0.1 (dashed black), 2 (dark). (D, H) $\lambda_G^+ = 0.006$ (light), 0.1 (dashed black), 2 (dark). Dashed line and number of realizations as in Fig. 2E-F of the main text.

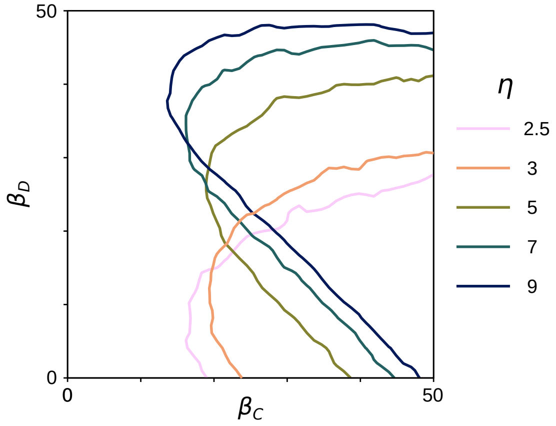


**Fig. S4**. Changes to the sync region boundary of the monomer binding scenario map in Fig. 5B of the main text as a function of the Hill exponent $\eta$, which controls the nonlinearity of the regulatory function. $\eta = 2.5$ corresponds to Fig. 5B of the main text and $\eta = 7$ to Fig. 5C. Sync region boundaries and number of realizations as in Fig. 2E-F of the main text.

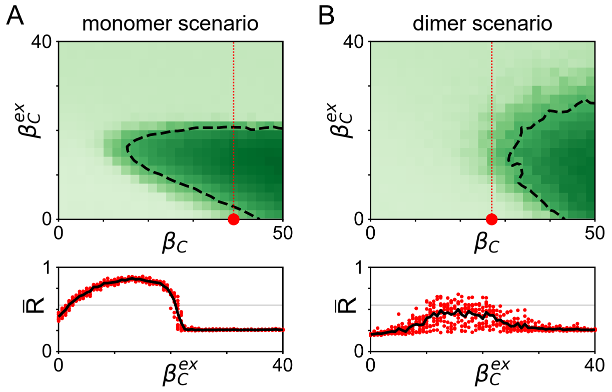


**Fig. S5**. Exogenous DeltaC expression assay in DeltaD mutant predicts distinct outcomes for monomer and dimer scenarios. We included an additional synthesis term in Eq. (52), $c_i' = ... + \beta_C^{ex}$, where $\beta_C^{ex}$ is an exogenous constant synthesis rate of DeltaC. Top: Steady state order parameter $\bar{R}$ in terms of endogenous and exogenous DeltaC synthesis rates $\beta_C$ and $\beta_C^{ex}$ for the (A) monomer and (B) dimer binding scenarios. Red dot is the DeltaD mutant condition for $\beta_C^{ex} = 0$ and vertical red dotted line indicates the cut plotted in bottom panels. Dashed line, color scale and number of realizations as in Fig.2C, D of the main text. Bottom: Steady state order parameter $\bar{R}$ as a function of $\beta_C^{ex}$ along the red dotted line in top panels for (A) monomer and (B) dimer binding scenarios. Red dots are 10 independent realizations and black line is the average. Horizontal gray line indicates the threshold $R_T$. Parameters as in corresponding scenario, except for $\beta_D = 0$, Table I.
